# Premature cleavage and polyadenylation in the minor intron of PTEN modulate its expression and generates a functional long non-coding RNA in breast cancer

**DOI:** 10.1101/2024.08.15.608023

**Authors:** Mariam Elesnawy, Rim Elghandour, Hana Hasna, Aisha Fakhroo, Boshra Al-Sulaiti, Ihab Younis

## Abstract

Minor introns constitute 0.4% of all introns in human cells, but they are unique in their ability to regulate the genes in which they are embedded such that their low splicing efficiency can be a rate limiting step in gene expression. The first intron of the tumor suppressor gene, PTEN, has been documented to be a minor intron, but very little is known about its regulation. Regulation of PTEN levels in cancer cells, especially breast cancer is tightly controlled as a very small reduction in its expression can lead to tumorigenesis. Indeed, many genetic, epigenetic, post-transcriptional and post-translational mechanisms are employed to reduce PTEN levels or activities, leading to cancer. Here, we uncover a previously unexplored mechanism for modulating PTEN expression utilizing its minor intron. The minor intron of PTEN minor intron has one of the lowest splicing efficacies in breast cancer cells, causing at least 50% of PTEN pre-mRNA to retain it, and thus not produce a functional protein. We also show that, unlike other minor introns, the retained intron in PTEN pre-mRNA is not used as a molecular switch that would be spliced out when needed but is rather processed by premature cleavage and polyadenylation into a long noncoding RNA (lncRNA) that we termed PINC. Exogenous expression of PINC caused significant alterations to cellular proliferation, including breast cancer cells that harbor a genetic mutation in the PTEN gene and do not produce a functional PTEN protein. This shows that this novel lncRNA functions independently of the encoded protein of the host gene.

## Introduction

Most genes in eukaryotes consist of exons that contain the information for protein expression, which are interrupted by relatively larger intervening sequences called introns. Since both exons and introns are transcribed into pre-mRNAs, it is necessary that the introns are spliced out and the exons joined together to form mature mRNAs (Wilkinson et al., 2020). Alternative splicing is a physiological process that generates different isoforms from the same pre-mRNA, leading to the production of different versions of the encoded protein of any given gene (Ast, 2004; Castle et al., 2008; Ellis et al., 2012; Kalsotra & Cooper, 2011; Nilsen & Graveley, 2010; Pan et al., 2008; Tapial et al., 2017; Weatheritt et al., 2016). Conversely, the disruption of normal splicing has been shown to lead to a large array of diseases, including cancer (El Marabti & Younis, 2018; Lee & Abdel-Wahab, 2016; Read & Natrajan, 2018). Aberrant splicing in cancer does not only affect tumor suppressor genes and oncogenes, but splicing deregulation is linked to all hallmarks of cancer, driving tumor progression and metastasis (Chabot & Shkreta, 2016; David & Manley, 2010; El Marabti & Younis, 2018; Fackenthal & Godley, 2008; Scotti & Swanson, 2016). While most introns are spliced out by the canonical spliceosome, around 0.4% of human introns (∼800 out of >200,000 introns) utilize a specialized spliceosome and are termed U12 introns or minor introns (mi-INTs) due to their low abundance (El Marabti et al., 2021). Minor introns are unique, as they are highly conserved and are non-randomly embedded in genes that play essential cellular processes (Olthof et al., 2019). They act as molecular switches as a post-transcriptional mechanism to promptly express hundreds of genes when needed (Younis et al., 2013). Deregulation of mi-INTs splicing could be detrimental to health, resulting in a wide range of ailments (El Marabti et al., 2021; Merico et al., 2015; Olthof et al., 2019; Younis et al., 2013).

Interestingly, the first of nine introns of the widely studied tumor suppressor gene (TSG), phosphatase and tensin homolog, *PTEN*, is a minor intron that we have previously shown to be regulated in response to cellular stress (Younis et al., 2013). PTEN functions in cell cycle regulation, metastasis, and apoptosis by downregulating the PI3K/AKT pathway (Cantley & Neel, 1999; Li et al., 1997; Steck et al., 1997; Tamura et al., 1998; Weng et al., 2001). Reduced expression of PTEN has been linked to several types of cancers with the mammary gland being the most sensitive (Garcia-Cao et al., 2012; Tamguney & Stokoe, 2007). The *PTEN* gene is a highly mutated locus in sporadic human tumors (Salmena et al., 2008). Given its importance in regulating cell growth and apoptosis, PTEN levels are highly regulated in cells, and even a subtle impairment can lead to tumorigenesis (Alimonti et al., 2010). In fact, *PTEN* exemplifies the continuum model of tumor suppressor genes (Alimonti et al., 2010; Berger et al., 2011; Malaney et al., 2017). This model suggests that complete loss of PTEN protein is not always required for tumor progression and a reduction as low as 20% can lead to tumor formation in lymph nodes and mammary glands (Alimonti et al., 2010), whereas 50% or 80% of PTEN protein is not enough to suppress tumors in mice that are hemizygous or hypomorphic, respectively (Gray et al., 1998; Trotman et al., 2003; Vidotto et al., 2023; Whang et al., 1998). Thus, cancer cells have developed several mechanisms to reduce PTEN levels either slightly or drastically. These include genetic alterations (e.g. point mutations, truncating mutations, and deletions), transcriptional inhibition (e.g. by modulating levels of transcription factors), epigenetic silencing, post-transcriptional repression (e.g. by inhibitory miRNAs), and post-translational modifications (e.g. phosphorylation, ubiquitylation, acetylation and oxidation), which alter protein level, function, localization, and interactions (Chow et al., 2007; Georgescu et al., 1999; Hettinger et al., 2007; Kwon et al., 2004; Li et al., 1997; Maccario et al., 2007; Meng et al., 2006; Okumura et al., 2006; Salmena et al., 2008; Virolle et al., 2001; Wu et al., 2003). In certain cancers, mutations that cause partial or complete loss of PTEN account for 17-46% of cases (Abida et al., 2019; Fusco et al., 2020; Grasso et al., 2012), however, in other cancers reduced PTEN levels are not readily explained by the above mechanisms, suggesting an unexplored process for inactivating this important tumor suppressor.

Given that (1) subtle reduction in PTEN can lead to tumor formation, (2) aberrant splicing can lead to protein loss associated with tumorigenesis (Lee & Abdel-Wahab, 2016), and (3) miINTs are known to be bottlenecks for expression of their host genes (Younis et al., 2013), we hypothesize that the regulation of PTEN pre-mRNA splicing, specifically its minor intron, presents a novel mechanism by which tumor cells alter PTEN expression. In this study, we used breast cancer cell lines to show that PTEN expression is indeed dependent on its minor intron splicing. Interestingly, we find that in breast cancer cells, the low efficiency of PTEN minor intron splicing and the consequential retention of the minor intron led to premature cleavage and polyadenylation in this intron, producing a previously unidentified PTEN lncRNA that functions independently of the PTEN protein in regulating breast cancer cell proliferation.

## Results

### PTEN minor intron splicing is altered in breast cancer cells

As a key tumor suppressor, it is expected that PTEN levels and/or function would be reduced in breast cancer cells. However, the known mechanisms for down-regulating PTEN, such as genetic alterations, transcriptional inhibition, epigenetic silencing, and post-transcriptional and post-translational modifications (Salmena et al., 2008) do not account for all breast cancer cases, suggesting an unknown mechanism for reducing PTEN in breast cancer cells. Given that the *PTEN* gene harbors a minor intron, which could be a rate limiting factor for mRNA production, we tested whether PTEN’s minor intron splicing efficiency is reduced in breast cancer, and whether this would affect PTEN protein expression. RNA sequencing (RNA-seq) data for MCF-7 (a breast adenocarcinoma) and MCF-10A (non-cancer breast cells) were downloaded from NCBI GEO (GSE71862) and aligned. Splicing efficiency in genes that harbor minor introns was then analyzed (unpublished data). Among the top affected genes, we found that the splicing of *PTEN’s* intron 1 in MCF-7 is significantly less efficient than that of MCF-10A (**Figure 1A**). As the *PTEN* minor intron is the first intron, it is expected that intron retention would affect the open reading frame, leading to reduced expression of the encoded protein. Thus, in breast cancer cells, two isoforms could be formed from PTEN pre-mRNA, where isoform 1 is fully spliced and does not include intron 1, hence can produce functional PTEN protein, while isoform 2 has minor intron retention, which leads to no PTEN protein being formed (**Figure 1B**). RNA-seq data were verified using RT-PCR from 3 different breast cancer cell lines (MDA-MB-231, MDA-MB-468, and MCF-7) (**Figure 1C**), confirming that intron 1 was retained at a remarkably high rate as indicated by the bands for unspliced PTEN. This was quantified using RT-qPCR, which shows that all three cell lines have more unspliced than spliced PTEN (**Figure 1D**). The levels of spliced PTEN are consistent, confirming that a large portion of what is being transcribed does not go on to be translated into a functional protein. Finally, western blot analysis showed that PTEN protein is indeed expressed from these cell lines from the properly spliced PTEN mRNA (**Figure 1E**). Of Note, MDA-MB-468 cells show no expression of PTEN protein due to a known nonsense mutation (De Vivo et al., 2000; Gasparyan et al., 2020).

**Figure 1:**
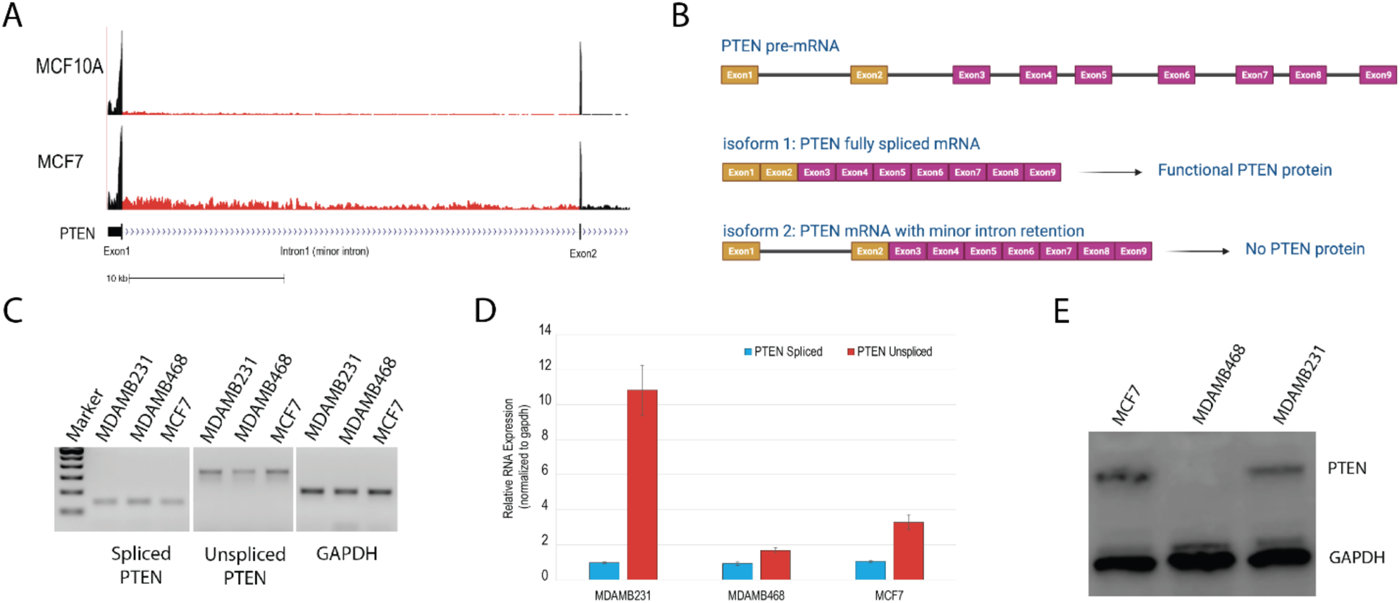
PTEN minor intron splicing is altered in breast cancer cells. (A) RNA-Seq data for MCF10A and MCF7 from NCBI GEO (GSE71862) showing expression of PTEN intron 1. (B) Schematic of PTEN minor intron splicing showing the two isoforms PTEN pre-mRNA can form to result in functional PTEN protein or no PTEN protein with minor intron retention. (C) Representative (N = 3) RT-PCR for spliced and unspliced PTEN on MDA-MB-231, MDA-MB-468, and MCF7 cell lines. GAPDH was used as a loading control. (D) Representative qPCR for spliced and unspliced PTEN on MDA-MB-231, MDA-MB-468, and MCF7 cell lines. Data shows the relative RNA expression after normalization to the loading control, GAPDH. Error bars represent standard deviations from three experiments. (E) Representative western blot showing PTEN and GAPDH (loading control) protein expression in MDA-MB-231, MDA-MB-468, and MCF7 cell lines.

### Inhibiting PTEN minor intron splicing affects cell viability

Our data suggests that PTEN’s minor intron retention is a mechanism that reduces the expression of PTEN in breast cancer cells, leading to inefficient tumor suppression. To assess these effects and confirm that PTEN’s minor intron is indeed a molecular switch, we designed antisense morpholino oligonucleotides (AMOs) to specifically bind exon1/intron1 junction and thus selectively inhibit the splicing of PTEN’s minor intron to only produce isoform 2 (**Figure 2A**). The efficiency of the AMOs was verified in all three cell lines, showing that 5µM PTEN AMO almost completely inhibited intron 1 splicing, as evidenced by the drastic decrease in the splicing indices (**Figure 2B**). Western blot analysis confirmed that PTEN protein was reduced by more than 87% and 95% upon transfecting cells with PTEN AMO in MDA-MB-231 and MCF7, respectively (**Figure 2C**). This data confirms that modulating the splicing of PTEN’s minor intron directly affects the levels of encoded protein.

**Figure 2:**
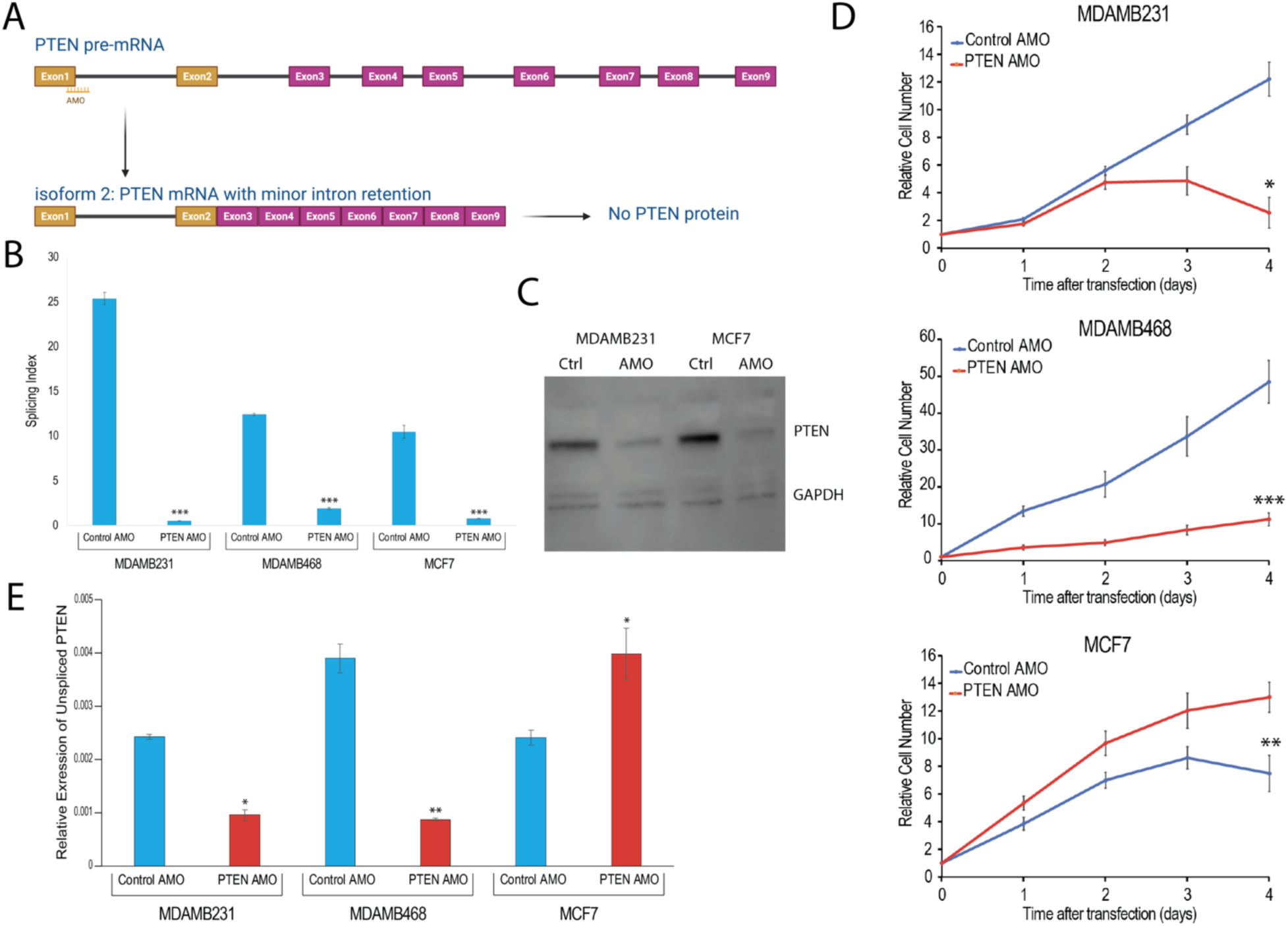
Inhibiting PTEN minor intron splicing affects cell viability. (A) Schematic of PTEN AMO that inhibits minor intron splicing resulting in the generation of isoform 2 that does not encode PTEN protein. (B) Representative qPCR for PTEN splicing index (spliced PTEN/unspliced + spliced PTEN) in cells transfected with control and PTEN AMO. (C) Western blot showing PTEN and GAPDH (loading control) protein expression in cells transfected with control and PTEN AMO. (D) RealTime-Glo™ MT Cell Viability Assay results represented as relative cell number for cells transfected with control and PTEN AMO. Results are normalized to day 0 and are for up to 4 days post transfection. For each condition, 10 wells were used as replicates, and error bars represent standard deviations. (E) Representative qPCR for relative unspliced PTEN expression in cells transfected with control and PTEN AMO. Statistical significance was measured using t-test except for proliferation data, in which a non-parametric (Mann-Whitney test) was used. *, **, and *** indicate p values < 0.5, 0.05, or 0.005, respectively. Error bars represent standard deviations from three experiments, unless otherwise indicated.

To confirm that modulating PTEN protein levels via minor intron splicing is consequential on cancer cell phenotype, cell proliferation was measured after transfecting cells with PTEN AMO. Surprisingly, selectively targeting PTEN minor intron splicing had a drastically different effect in the triple negative cell lines (MDA-MB-231 and MDA-MB-468), compared to the hormone responsive cell line (MCF7) (**Figure 2D**). As PTEN AMO inhibits the splicing of intron 1 of PTEN, leading to reduced translation of a functional tumor suppressor protein, we expected that it would increase cell proliferation. While this was the observed effect in MCF7 cells, PTEN AMO treatment caused an unexpected reduction in growth of MDA-MB-231 and MDA-MB-468 (**Figure 2D**). This was particularly surprising in MDA-MB-468 cells that do not express a functional PTEN protein.

Upon further inspection, we found a strong correlation between the effect on proliferation and the levels of unspliced PTEN produced after PTEN AMO transfection (**Figure 2E**). In MDA-MB-231 and MDA-MB-468, PTEN AMO did not only affect spliced PTEN expression, but the expression of unspliced PTEN decreased as well, suggesting that the transcripts with a retained intron are unstable in these cell lines. Alternatively, transfection of MCF7 with PTEN AMO causes the expected decrease in spliced PTEN (**Figure 2B**), but the retained unspliced intron accumulated, which seems to correlate with an increase in proliferation. We thus suggest that it is the unspliced PTEN that is needed to sustain proliferation, and that changes in unspliced PTEN are somehow dominant over changes in spliced PTEN and PTEN protein. Taken together, these data suggest that while the inhibition of PTEN’s minor intron does cause the expected reduction in PTEN protein, it leads to more tumor suppressive activity in some triple negative cell lines, which corresponds with the changes in unspliced PTEN.

### PTEN inhibition affects cell viability in a cell line-dependent manner

To distinguish between the effect of altering the levels of the spliced mRNA, unspliced mRNA or PTEN protein on cancer cell proliferation, we utilized two additional approaches: a PTEN protein activity inhibitor (SF1670) and PTEN siRNA. We focused on MDA-MB-231 and MDA-MB-468 because these triple negative cell lines showed unexpected results with the PTEN AMO. In addition, MDA-MB468 does not produce a functional PTEN protein, making it a good control for dissecting these effects.

SF1670 is a pharmacological inhibitor that binds to PTEN protein and inhibits its function without affecting the protein levels, which was what we observed (**Figure 3A**). Conversely, PTEN siRNA is expected to reduce the expression of both PTEN mRNA and protein. Indeed, PTEN mRNA was reduced by 66-69% (**Figure 3B**) and the protein was decreased by 71% after PTEN silencing (**Figure 3C**). In both cases, PTEN inhibition should presumably result in increased cell proliferation as the inhibition of PTEN protein activity or its levels should decrease its tumor suppressive activity. However, this effect should be limited to MDAMB231 cells because they express functional PTEN protein. Indeed, both SF1670 and PTEN siRNA resulted in increased cell proliferation in MDA-MB-231 whereas MDA-MB-468 showed little to no effect (**Figures 3D and 3E**). This data suggests that these cells respond as expected when PTEN mRNA or protein are targeted via the traditional methods. As such, the unexpected effects of the PTEN AMO in the triple negative cell lines seem to be unique to the AMO’s ability to modulate unspliced PTEN mRNA rather than the protein levels.

**Figure 3:**
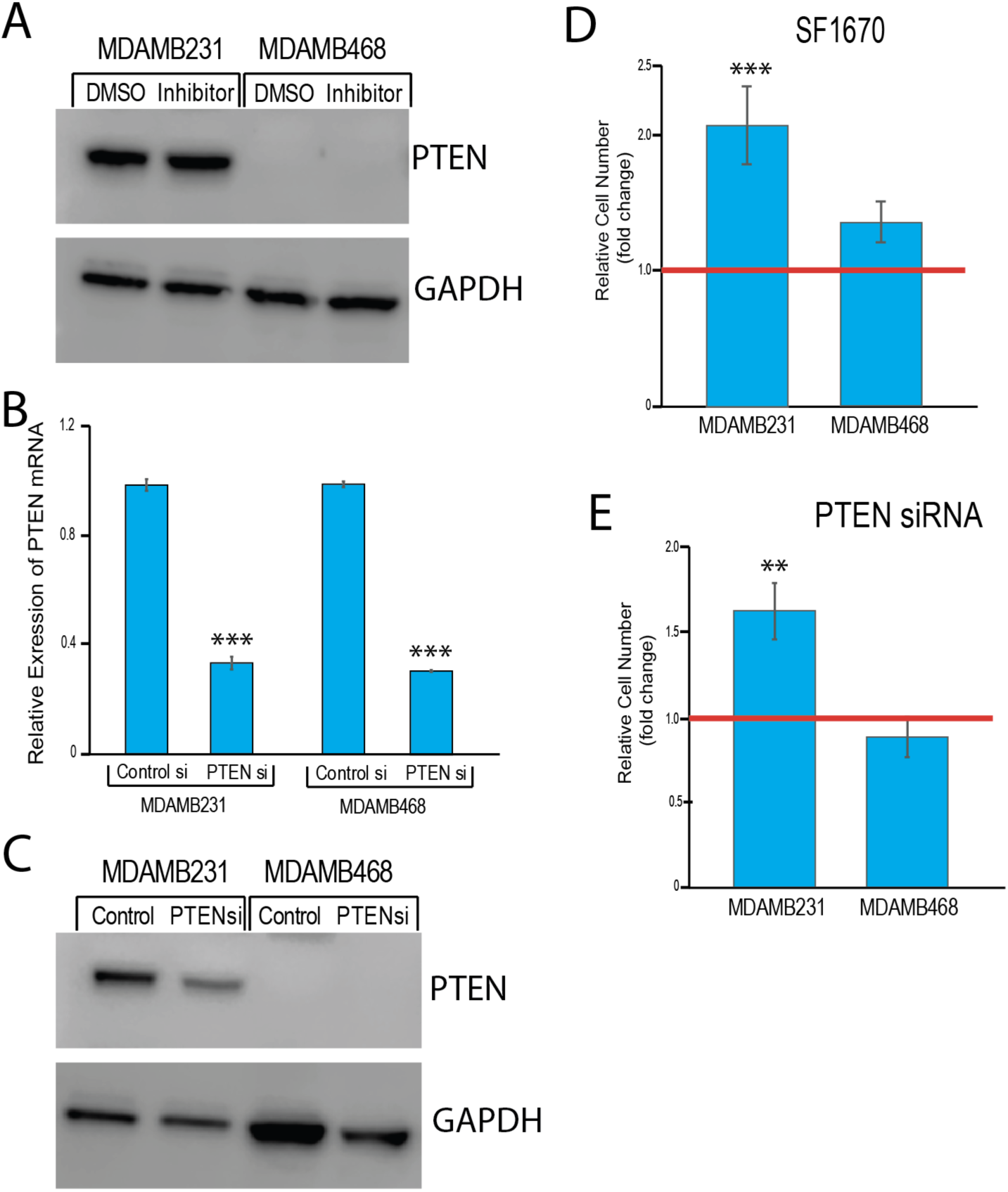
PTEN inhibition affects cell viability in a cell line-dependent manner. (A) Western blot showing PTEN and GAPDH (loading control) protein expression in cell lines treated with DMSO and SF1670. (B) Representative qPCR for relative PTEN mRNA expression in cell lines transfected with control and PTEN siRNA. (C) Western blot showing PTEN and GAPDH (loading control) protein expression in cell lines transfected with control and PTEN siRNA. (D and E) RealTime-Glo™ MT Cell Viability Assay results represented as relative cell number (fold change) for cell lines treated with DMSO and SF1670 (Figure D) or transfected with control and PTEN siRNA (Figure E). Results Show relative cell number 48 hours after treatment and are normalized to day 0. For each condition, 10 wells were used as replicates, and error bars represent standard deviations. Statistical significance was measured using t-test except for proliferation data, in which a non-parametric (Mann-Whitney test) was used. *, **, and *** indicate p values < 0.5, 0.05, or 0.005, respectively. Error bars represent standard deviations from three experiments, unless otherwise indicated.

### A retained PTEN minor intron is processed into a poly-adenylated lncRNA

To investigate the discrepancy of the effects of inhibiting PTEN’s minor intron splicing using a PTEN AMO between different breast cancer cells, we further examined the unspliced PTEN mRNA. We first checked the Genotype-Tissue Expression (GTEx) Project for expression of PTEN and its minor intron splicing efficiency in various normal tissues. Interestingly, many tissues showed some level of intron 1 retention (**Figure 4A**). However, upon closer inspection, the intron signal did not conform to the typical intron retention, but was rather reminiscent of premature cleavage and polyadenylation (PCPA) that has been previously observed upon inhibition of U1 snRNP (Berg et al., 2012; Kaida et al., 2010). **Figure 4A** shows examples of 12 tissues with varying levels of detectable RNA-seq reads in intron 1 of PTEN. Remarkably all the tissues show an exact point of attrition at around 2500 bases into the intron. The signal of spliced exons (exon 1 and exon 2 shown in **Figure 4A**) does not seem to be affected, suggesting that in normal physiological conditions most of the pre-mRNA is spliced properly and goes on to produce a PTEN protein, whereas a small fraction of the pre-mRNA undergoes intronic premature cleavage and polyadenylation, does not form a full-length transcript, and goes on to produce a long noncoding RNA (lncRNA).

**Figure 4:**
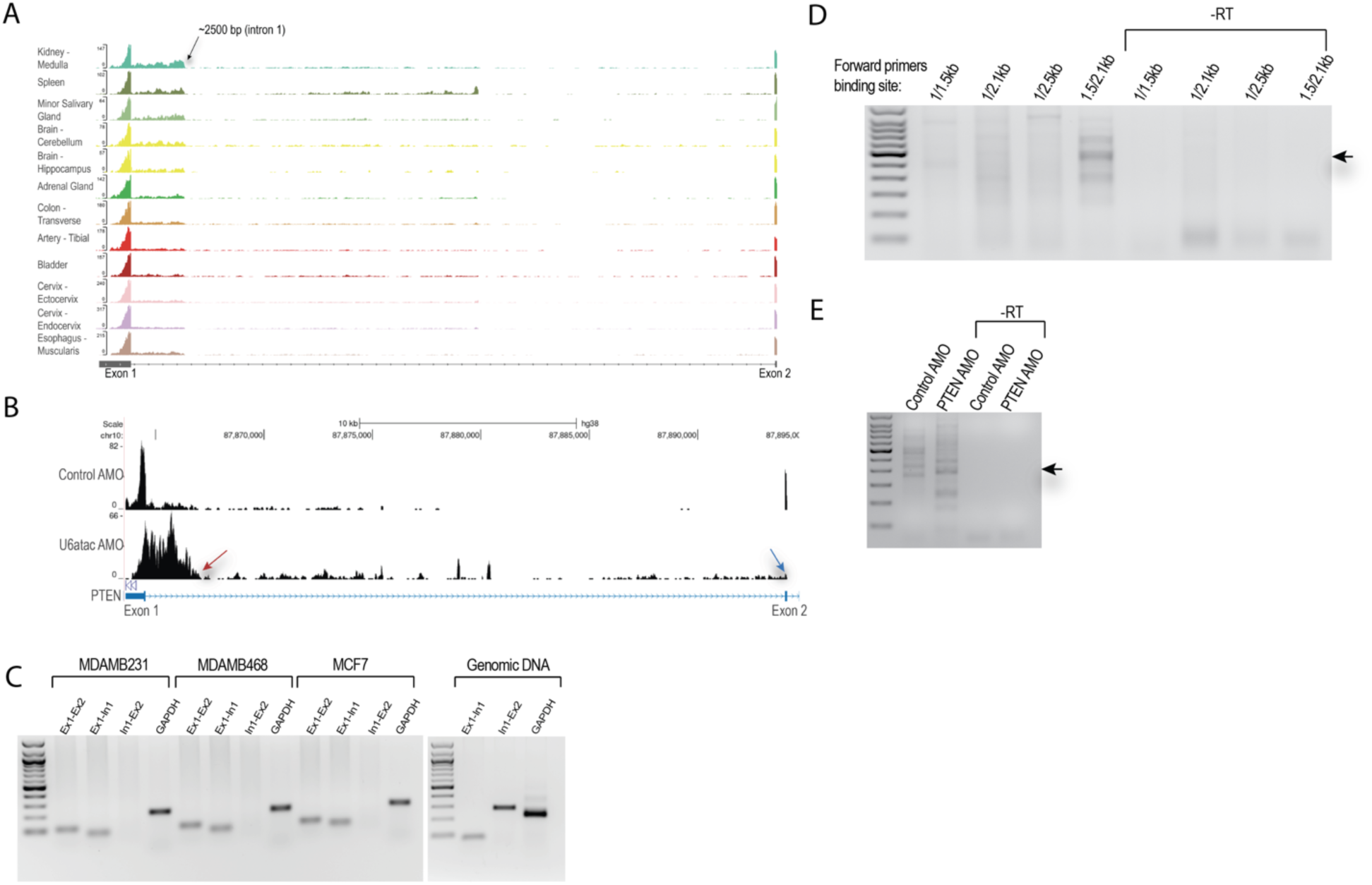
A retained PTEN minor intron is processed into a poly-adenylated lncRNA. (A) Expression of PTEN minor intron in various normal tissues. Data was extracted from the GTEx project. (B) RNA-Seq data following U6atac AMO transfection in MDA-MB-231 cells, showing PTEN expression in control and U6atac AMO conditions. (C) Representative RT-PCR for PTEN exon 1-exon 2 junction (spliced), exon 1-intron 1 junction (unspliced at 5’ end of intron 1), and intron 1-exon 2 junctions (unspliced at 3’ end of intron 1) in MDA-MB-231, MDA-MB-468, and MCF7 cell lines. GAPDH was used as a loading control and genomic DNA as a positive control. (D and E) Representative 3’ RACE on RNA extracted from cells transfected with U6atac AMO (Figure D) and control and PTEN AMO (Figure E) on different primers binding different regions in intron 1.

To confirm the enrichment of the polyadenylated lncRNA and rule out that our data are a side effect of PTEN AMO and its binding to the 5’ splice site of the PTEN pre-mRNA, we inhibited PTEN splicing through a different approach. We have previously used a U6atac AMO, which inhibits all minor intron splicing by masking the catalytic subunit of the minor spliceosome (Younis et al., 2013). As such, U6atac AMO does not interact with PTEN pre-mRNA and its effects are indirect. As shown in **Figure 4B**, transfecting MDA-MB-231 cells with U6atac AMO caused the exact pattern of PCPA at the same location (red arrow) that is observed in normal tissues. The difference here was the complete loss of Exon 2 signal (blue arrow) and the spliced Exon 1-Exon 2 junction reads, confirming that complete inhibition of the splicing of PTEN’s minor intron generates a lncRNA at the expense of spliced mRNA.

The data above suggests that PCPA took place within the first 20% of the intron and only the 5’ end of the processed transcript is stable. We thus designed primers that specifically amplify either the 5’ or the 3’ end of PTEN’s first intron. RT-PCR for RNA extracted from all three cell lines consistently showed the expected spliced PTEN as shown by the primer pair that detects Exon 1-Exon 2 junction (Ex1-Ex2) (**Figure 4C**). However, when the exon-intron boundaries were amplified, the primer pair that detects Exon 1-Intron 1 boundary (Ex1-Int1) showed a clear signal, whereas those primers that detect Intron 1-Exon 2 boundary (Int1-Ex2) showed no signal. We confirmed the ability of the primers (Int1-Ex2) to amplify the 3’ end of the intron using genomic DNA as input. These data strongly suggest that PTEN’s intron 1 is not fully retained as an intact intron but only the 5’ end of it accumulates in cells.

For PCPA to occur in PTEN’s minor intron, an intronic polyadenylation signal (iPAS) needs to be utilized. Consequently, the intron is cut and a polyA tail is added, generating a stable polyadenylated transcript. To test this, we used a polyA detector tool (https://apa.cs.washington.edu/detect) which identified several iPASs, including a strong one at position 2469 in the intron. This is highly consistent with the data observed in GTEx and U6atac AMO transfection. To verify the usage of these iPASs, 3’ RACE was employed on cells transfected with U6atac AMO. Utilizing several forward primers in the intron and oligodT reverse primers, we scanned for polyadenylated RNAs in the intron. This showed a clear band corresponding to a polyadenylated RNA at ∼2500 in PTEN’s intron 1 (**Figure 4D**), supporting our hypothesis that the retained intron 1 of PTEN is processed into a polyadenylated RNA. This was also detected in samples transfected with PTEN AMO (**Figure 4E**).

Checking the sequence of Exon 1 plus 2500 bases from Intron 1 did not show any significant open reading frame, suggesting that this polyadenylated RNA is a long non-coding RNA that we termed <u>P</u>TEN intronic <u>n</u>on-<u>c</u>oding (PINC) RNA.

### PINC RNA affects cell viability differently in various breast cancer cell lines

To check whether PINC RNA is a functional lncRNA, we cloned various fragments of PTEN’s Exon 1 plus Intron 1 into an expression vector and over-expressed them in breast cancer cell lines. The data for the different fragments are similar, so only the shortest functional fragment (termed PINC1) is shown here. PINC1 represents 79 bases from Exon 1 and 259 bases from Intron 1. Various cell lines were transfected with 250 ng of the expression plasmids (empty or containing a PINC fragment) for 24 hours. RNA and proteins were then collected to check for overexpression of the ncRNA and for any effects on the endogenous PTEN expression. We used 250 ng of plasmid to achieve an over-expression that is close to physiological conditions and to avoid non-specific toxicity effects. To measure PINC expression, the PTEN unspliced primers were used because data in Figure 4C confirmed that most, if not all, unspliced PTEN’s minor intron is processed into a lncRNA. This showed that PINC overexpression was 2.7, 2.4 and 2.1-fold above control in MDA-MB-231, MDA-MB-468 and MCF7, respectively (**Figure 5A**). While RT-PCR showed that PINC overexpression was successful but limited to less than 3-fold increase, it also showed that this amount of overexpression did not have any significant effects on endogenous spliced PTEN (**Figure 5A**). In addition, no effects were observed on the production of PTEN protein from the endogenous gene (**Figure 5B**). This suggests that PINC does not regulate the expression of its host gene. Interestingly, PINC overexpression caused varying effects on the proliferation of different breast cancer cell lines. As shown in **Figure 5C**, PINC overexpression caused a small decrease in MCF7 proliferation, but this was not statistically significant (repeated 3 times). Surprisingly, the two triple negative breast cancer cell lines responded in opposite ways to PINC overexpression. While MDA-MB-231 cells proliferated significantly more, the MDA-MB-468 cells proliferation was significantly decreased (**Figure 5C**). It is noteworthy that one of the main differences in these cells is the lack of PTEN protein production in MDA-MB-468. Taken together, the data suggests that PINC has a role in regulating cellular proliferation, and this role might be affected by the presence or absence of a functional PTEN protein in cells.

**Figure 5:**
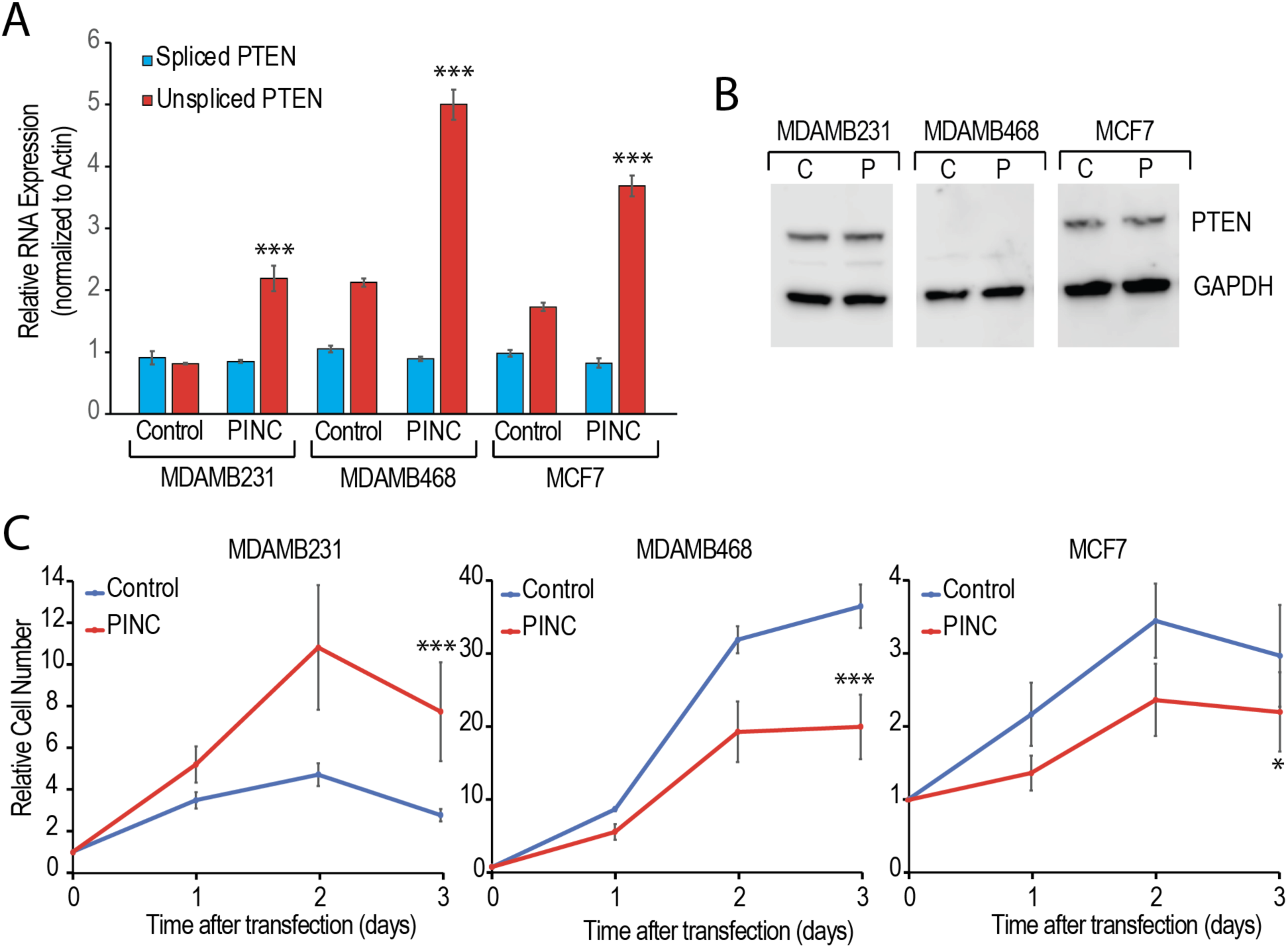
PINC RNA affects cell viability differently in various breast cancer cell lines. (A) Representative qPCR for spliced and unspliced PTEN on MDA-MB-231, MDA-MB-468, and MCF7 cell lines transfected with pcDNA3.1A (control) and PINC RNA-expressing plasmid. Data shows the relative RNA expression after normalization to the loading control, actin. Error bars represent standard deviations from three experiments. (B) Western blot showing PTEN and GAPDH (loading control) protein expression in MDA-MB-231, MDA-MB-468 and MCF7 cell lines transfected with control (C) and PINC (P) plasmids. (C) Cell viability using RealTime-Glo™ MT Cell Viability Assay results represented as relative cell number for MDA-MB-231, MDA-MB-468, and MCF7 cell lines transfected with control and PINC plasmids. Results are normalized to day 0 and are for up to 3 days post transfection. For each condition, 10 wells were used as replicates, and error bars represent standard deviations. Statistical significance was measured using t-test except for proliferation data, in which a non-parametric (Mann-Whitney test) was used. *, **, and *** indicate p values < 0.5, 0.05, or 0.005, respectively. Error bars represent standard deviations from three experiments, unless otherwise indicated.

## Discussion

As a tumor suppressor, PTEN plays a pivotal role in regulating cellular proliferation, with even a small decrease in its levels leading to tumor formation in some tissues, including breast tissue. As such, many mechanisms are employed to ensure a tight regulation of PTEN expression and its activity in normal tissues. Deregulating any of these mechanisms can thus lead to cancer. While genomic alterations such as point mutations, truncating mutations, and deletions are well documented, other non-genetic aberrations can reduce PTEN’s expression and/or activity. These include transcriptional inhibition, epigenetic silencing by promoter methylation, post-transcriptional repression by inhibitory miRNAs, and post-translational modifications (Chow et al., 2007; Georgescu et al., 1999; Hettinger et al., 2007; Kwon et al., 2004; Li et al., 1997; Maccario et al., 2007; Meng et al., 2006; Okumura et al., 2006; Salmena et al., 2008; Virolle et al., 2001; Wu et al., 2003). In this study, we focused on a less explored mechanism: alternative splicing. An early study by Sarquis et al showed an interesting correlation between PTEN’s differential expression, its splice variants (SV), and the pathogenesis of breast cancer. Different SVs have an effect on downstream elements of the AKT pathway, such as Cyclin D1. For example, increased SV5b protein expression enhances cyclin D1 promoter activity, opposite to PTEN’s function (Sarquis et al., 2006). On the other hand, Agrawal and Eng (2006) demonstrated that SV5b is overexpressed in sporadic breast cancer patients, while in Cowden Syndrome patients, SV3a is overexpressed, suggesting that the differential expression of PTEN SVs may be the cutoff between some benign and cancerous tumors (Agrawal et al., 2005; Agrawal & Eng, 2006). While these reports establish the importance of alternative splicing in regulating PTEN’s activity, they all focus on alternative exon splicing. As PTEN’s first intron is a minor intron and we have demonstrated that it is inefficiently spliced in cells, we investigated here the regulation of intron retention and its correlation with PTEN expression and activity. Moreover, we have previously shown that PTEN expression can be regulated by increasing the efficiency of its minor intron splicing in a p38 MAPK-dependent manner in cervical cancer cells (Younis et al., 2013). Thus, we hypothesized that deregulation of PTEN’s minor intron splicing is a novel mechanism to limit PTEN expression in cancer.

Our data confirms that in several breast cancer cell lines, the first intron in a large fraction of PTEN’s pre-mRNA is not spliced efficiently producing an isoform that cannot be translated into a functional protein. The use of an antisense morpholino that blocks the 5’ splice site of intron 1 resulted in an almost complete loss of PTEN spliced mRNA, and consequently the encoded protein, but resulted in unexpected effects on cellular proliferation. While MCF7 cells responded as expected when a tumor suppressor such as PTEN is reduced with increased proliferation, the two triple negative cell lines used in this study showed drastic decrease in cellular proliferation. This is intriguing as the PTEN spliced mRNA and protein were both reduced by more than 90%, suggesting an almost complete loss of tumor suppression. However, the cells grew less than the control. A more surprising result was the response of MDA-MB-468 cells, where we observed a decrease in cell survival with PTEN AMO. MDA-MB-468 cells harbor a nonsense mutation in the *PTEN* gene that leads to complete loss of the protein (De Vivo et al., 2000; Gasparyan et al., 2020). Thus, it is not at all expected that MDA-MB-468 cells would respond to reduction in PTEN expression, which was indeed the case when PTEN’s expression was reduced by siRNA or its activity by a chemical inhibitor. In both cases, the proliferation of MDA-MB-231 cells increased, whereas that of MDA-MB-468 did not significantly change. This led us to conclude that the effects of inhibiting the splicing of PTEN’s minor intron are at least partially independent of reducing the proteins in cells.

Upon further inspection of PTEN’s minor intron, we identified at least 12 tissues from GTEx in which the first 2500 bases of the intron are notably accumulated as opposed to the rest of the intron. We confirmed this experimentally in the breast cancer cell lines, showing a 100% biased accumulation of the 5’ end of the intron as opposed to the 3’ end when minor intron splicing was inhibited using U6atac AMO, which inhibits the splicing machinery without binding to the target pre-mRNAs. We opted to use this alternative approach to inhibit PTEN’s minor intron splicing to avoid side effects of direct AMO binding to PTEN’s minor intron and causing artificial stabilization of the pre-mRNA. Like the GTEx data, U6atac AMO resulted in accumulation of the first 2500 bases of the intron but was also accompanied by loss of signal in Exon 2. This strongly suggested that premature cleavage and polyadenylation occurred resulting in a previously unidentified transcript. Our data show that this transcript is polyadenylated and is unlikely to have an open reading frame. We thus termed it <u>P</u>TEN intronic <u>n</u>on-<u>c</u>oding (PINC) RNA. Remarkably, when PINC RNA was exogenously expressed from a plasmid, it caused different effects on proliferation in the triple negative cell lines without affecting the endogenous expression of PTEN’s spliced mRNA or protein. Again, MDA-MB-468 cells provided an intriguing case due to their lack of PTEN protein. These cells proliferated less when PINC RNA was exogenously expressed, as opposed to MDA-MB-231, which proliferated more. It was surprising that the effects of exogenous expression of PINC RNA were different than producing it from the endogenous pre-mRNA using PTEN AMO. While MDA-MB-468 had similar effects (reduced proliferation) with either treatment, MCF7 and MDA-MB-231 cell lines had opposite effects. We speculate that the effects on cellular proliferation are due to the balance of many factors, some of which have opposite functions. These include the presence or absence of PTEN protein, the absolute levels of spliced mRNA and unspliced mRNA (PINC RNA), the levels of downstream effectors such as miRNAs and RNA binding proteins (see next section) and other factors that are expected to be different when an RNA is expressed exogenously vs. endogenously.

We classified PINC RNA as a long noncoding RNA. lncRNAs are typically similar to protein-coding mRNAs in terms of their transcription regulation and post-transcriptional processing (5′ end capping, 3′ end polyadenylation and alternative splicing) (Kopp & Mendell, 2018), but they are less conserved, are expressed at lower levels than mRNAs, and tend to have more tissue-specific expression (Derrien et al., 2012; Djebali et al., 2012; Fazal et al., 2019; Kopp & Mendell, 2018). There are many reasons for why a lncRNA would have different effects in different cell lines. Two of the most studied mechanisms by which lncRNAs function in cells, including cancer, are sequestering and thus inhibiting miRNAs and RNA binding proteins (RBPs) (Jonas et al., 2020; Ratti et al., 2020; Shaath et al., 2022; X. Zhang et al., 2019). Given that the expression profiles of miRNAs and RBPs are different in the cell lines studied, it is reasonable to speculate that potential targets might be missing or over-represented in one cell line vs. another. Thus, the effect of sequestering them would be different. In this study, we used the smallest functional fragment (∼340 bases) of PINC RNA to allow us to identify potential miRNAs and RBPs that might be sequestered by this RNA. Indeed, we identified hsa-miR-542-3P and hnRNP D as the top candidates for binding to PINC RNA. Sequestering hsa-miR-542-3P or hnRNP D, or a combination of both (and potentially other differentially expressed targets) would be expected to affect cellular proliferation as both of them have been previously shown to play a role in regulating cancer cell behavior (Althoff et al., 2015; Al-Tweigeri et al., 2022; Fan et al., n.d.; Gouble et al., 2002; He et al., 2014; Yoon et al., 2010). Further detailed work is needed to delineate the exact mechanism by which PINC RNA regulates cellular proliferation, including its sequestration of hsa-miR-542-3P and hnRNP D.

It was surprising that our study unmasked a previously unidentified mechanism for regulating a gene as studied as PTEN. On the other hand, PINC RNA expression could have been overlooked as inefficient splicing, which is common in cancer (Rahman et al., 2020; J. Zhang & Manley, 2013). In addition, it was likely overlooked as high PINC RNA production is dependent on both intronic premature cleavage and polyadenylation (iPCPA) and alternative splicing, specifically intron retention. While alternative cleavage and polyadenylation towards the 3’ end of pre-mRNAs is a prevalent process and well documented (Dharmalingam et al., 2022; Xu et al., 2021; Zheng & Tian, 2014), iPCPA has recently emerged as a significant mechanism to regulate gene expression (Berg et al., 2012; Kaida et al., 2010). Conversely, there are only a few documented cases of alternative splicing leading to the production of lncRNAs from what is typically a protein-coding gene. These include the PNUTS and PD-L1 protein coding genes that can be alternatively spliced to produce lncRNAs that promote tumor progression (Grelet et al., 2017; Liu et al., 2023; Qu et al., 2021).

This study provides the first evidence of a PTEN intronic ncRNA (PINC) that is produced when the first intron of PTEN is inefficiently spliced, followed by intronic cleavage and polyadenylation. Overexpressing PINC RNA in breast cancer cells has significant effects on cellular proliferation. While the mechanisms by which PINC RNA regulates proliferation are still poorly understood, they are unlikely to be through directly regulating PTEN’s own mRNA and/or protein production. Although we cannot rule out a functional relationship between PTEN activity and PINC RNA, it would be of paramount importance to further understand this relationship as regulating PTEN levels or activity can have profound therapeutic effects especially in breast cancer.

## Materials and Methods

### Cell Culture

Human breast cancer cell lines MDA-MB-231, MDA-MB-468 and MCF-7 purchased from ATCC were maintained in Dulbecco’s modified Eagle’s Medium (DMEM) with high glucose, GlutaMAX™, supplemented with 10% Fetal Bovine Serum (FBS), 1% (100U/ml) Penicillin-Streptomycin mixture, and 1% non-essential amino acids. Cells were grown at 37°C in a 5% CO_2_ humidified incubator. At 70-80% confluence, cells were passaged.

### Transfection with AMOs

Control (5′-CCT CTT ACC TCA GTT ACA ATT TAT A-3′), PTEN (5’-CGC AGA AAT GGA TAC AGG TCA AGT C-3’) and U6atac (5′-AAC CTT CTC TCC TTT CAT ACA ACA C-3′) antisense morpholino oligonucleotides (AMOs) were synthesized by GeneTools, and up to 5 μM were transfected into 1 million cells using electroporation by Neon 100uL Transfection system (Invitrogen). Cells were resuspended in resuspension buffer R, provided with the transfection kit. The AMOs were then added to the cells and electroporated, according to the manufacturer’s protocol. Transfected cells were then plated in complete medium lacking Penicillin-Streptomycin. RNA and proteins were harvested from transfected cells one day post transfection. Transfected cells were also plated for cell proliferation.

### PINC Plasmid Design and Transfection

To clone the smallest functional fragment of PTEN’s intronic noncoding RNA or PINC1, 79 bp of exon 1 plus 259 bp of intron 1 were amplified by PCR using primers in Table 1 and cloned into pcDNA3.1A. Successful inserts were confirmed using sequencing. For exogenous expression, 250ng of an empty plasmid (control) or a plasmid expressing PINC1 were transfected into 50,000 breast cancer cells using Lipofectamine LTX reagent according to manufacturer’s recommendation. Total RNA and proteins were extracted 24 hours post-transfection as described below.

**Table 1:**
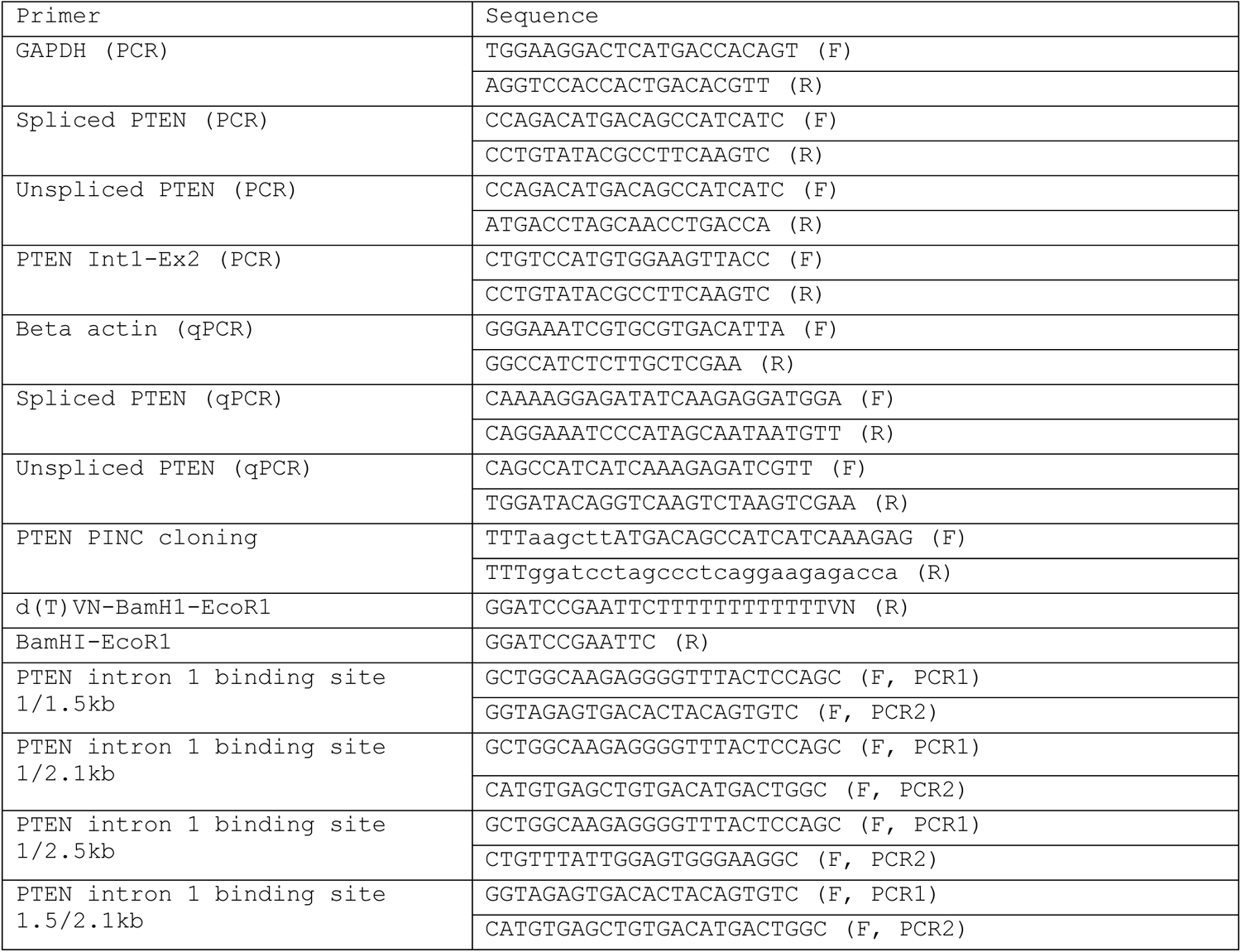
Primer Sequences.

### Cell Proliferation Assay

Cell proliferation was evaluated using RealTime-Glo™ MT Cell Viability Assay (Promega). After transfection, cells were plated at a density of 2000-3000 cells/well in an opaque-walled 96-well plate. For each treatment, 10 wells were used as replicates. The MT Cell Viability Substrate and NanoLuc Enzyme were heated to 37°C, diluted in complete media and added at a concentration of 1X for a total volume of 100μL per well. Cells were then incubated for 30 minutes at 37°C. Luminescence was measured on a GloMax® Explorer Multimode Microplate Reader (Promega) at 37°C, with an integration time of 0.3 seconds per well. Measurements were taken at the time of plating (Day 0), and subsequently on a daily basis for 3-4 days.

### PTEN inhibitor treatment

MDA-MB-231 and MCF-7 cells were plated at a density of 500,000 cells/well in 6-well plates and allowed to attach for 24 hours. SF1670 was purchased from Sigma-Aldrich (SML0684-5MG), and cells were treated with 0.3uM SF1670, diluted in dimethyl sulfoxide (DMSO). After 48 hours of incubation, RNA and protein were extracted as described below. Cellular proliferation was also measured at Day 0 and Day 2 using RealTime-Glo™ MT Cell Viability Assay (Promega).

### RNA and Protein Extraction

RNA was extracted from cells following transfection or inhibitor treatments using Qiagen’s RNeasy Plus Mini Kit (Catalog #: 74136), according to manufacturer’s recommendations. Protein was harvested from live cells using RIPA buffer with protease and phosphatase inhibitor cocktail. Protein concentrations were measured using the Bio-Rad protein assay.

### RT-PCR and qPCR

For reverse transcription, 1 μg of RNA was converted to cDNA using the ProtoScript® II First Strand cDNA Synthesis Kit (New England BioLabs), according to the manufacturer’s protocol. cDNA was then diluted 10-fold in nuclease-free water and used as input for PCR. Primers sequences are in table 1. PCR was carried out using OneTaq® 2X Master Mix kit (New England BioLabs), according to manufacturer’s recommendations. Cycling conditions were: denaturing of DNA at 94°C for 30 seconds; 32 cycles of 30 seconds at 94°C for denaturation, 30 seconds at 50°C for annealing, and 30 seconds at 68°C for extension; and a final extension at 68°C for 5 minutes. Amplified products were run on 2% TAE agarose gels and imaged in a UV box. For qPCR, expression of spliced and unspliced PTEN was quantified using either PowerUp (Applied Biosystems) or PowerTrack (Applied Biosystems) SYBR Green master mix. Spliced PTEN primers were designed to bind exon 1 (forward) and exon 2 (reverse), whereas unspliced PTEN primers were designed to bind exon 1 (forward) and exon 1-intron 1 junction (reverse). qPCR was performed on the QuantStudio 6 system (Applied Biosystems), and relative expressions were normalized to beta actin levels. Primer sequences are in Table 1.

### Western Blotting

Extracted total proteins were diluted in Tris-Glycine SDS buffer and β-mercaptoethanol, then denatured at 95°C for 5 minutes and loaded on 4-12% NuPAGE™ Bis-Tris gels (Invitrogen). Samples were transferred to a nitrocellulose membrane using a semi-wet transfer. The membrane was blocked with 5% BSA in TBST for 2 hours to overnight at room temperature, and then incubated with primary antibodies overnight at 4°C. The primary antibodies used were: rabbit monoclonal anti-PTEN antibody (Cell Signaling Technology) and mouse monoclonal anti-GAPDH antibody (Cell Signaling Technology). Following washes with TBST, the HRP-conjugated secondary antibodies were added at 1:5000 dilution and incubated for 1 hour at room temperature, followed by washes with TBST. Membrane was then imaged using the Li-Cor Odyssey® Fc Imager.

### RNA-Sequencing

Total RNA from MDA-MB-231 and MCF7 cells that were transfected with AMOs was extracted as described above. Total RNA with a RIN number above 8 was used as input for library preparation using TruSeq Stranded mRNA kit from Illumina following the manufacturer’s protocol. Briefly, from 500 ng of total RNA, mRNA molecules were purified using poly-T oligo attached magnetic beads and then mRNA was fragmented. cDNA was generated from the cleaved RNA fragments using random priming during first and second strand synthesis. Barcoded DNA adapters was ligated to both ends of the DNA and then amplified. The quality of library generated was checked on an Agilent 2100 Bioanalyzer system and quantified using Qubit system. Libraries that pass quality control were pooled and sequenced on a NextSeq2000 system at a minimum of 20 million paired-end reads (2x75 bp) per sample. Basic trimming and quality control were performed during the conversion of raw data to fastq using Illumina BCL2Fastq Conversion Software. For alignment and expression analysis, an R pipeline was used, including the following packages: RSubread, DESeq, and edgeR.

### PAS Detection and 3’ RACE

The polyA detector tool (https://apa.cs.washington.edu/detect) was used to identify PAS signals, where the PTEN minor intron sequence was inputted. For 3’ RACE, total RNA was extracted as described above. PolyA tail-specific primer oligo-d(T)VN-BamH1-EcoR1 (Table 1) was used to reverse transcribe poly-adenylated mRNA into cDNA using ProtoScript® II First Strand cDNA Synthesis Kit (New England BioLabs) as described above. The reverse-transcribed cDNA was then diluted 10-fold in nuclease-free water. The nested PCR included two rounds of amplification. For PCR1, several forward primers that hybridize to PTEN intron 1 were used (Table 1), while the reverse primer was the same polyA tail-specific primer oligo-d(T)VN-BamH1-EcoR1. Amplification conditions: 30 seconds at 94°C; 25 cycles of 30 seconds at 94°C, 30 seconds at 50°C, and 2 minutes at 68°C; and a final extension of 5 minutes at 68°C. For PCR 2, 5ul of the amplification product of PCR 1 was used as input, and amplified using one of the many forward primers that hybridize to intron 1 (Table 1), in addition to a reverse primer that corresponds to BamH1-EcoR1 sequence (Table 1), ensuring that only poly-adenylated mRNA that were amplified in PCR1 are a proper template for PCR2. For PCR2, the same conditions were used as PCR1 except for the number of cycles time, which was 35 in this case.

## Acknowledgements

We are grateful to the members of the Biological Sciences program, especially Maya Kemaldean for comments on the manuscript.

